# A unique and biocompatible approach for corneal collagen crosslinking in vivo

**DOI:** 10.1101/2024.03.18.585574

**Authors:** Ayesha Gulzar, Humeyra N Kaleli, Gulsum D Koseoglu, Murat Hasanreisoglu, Ayşe Yildiz, Afsun Sahin, Seda Kizilel

## Abstract

Corneal crosslinking (CXL) is a widely applied technique to halt the progression of ectatic diseases by increasing the thickness and mechanical stiffness of the cornea. This study investigated the biocompatibility and efficiency of a novel CXL procedure using ruthenium and blue light in rat corneas and evaluated factors important for clinical application. To perform the CXL procedure, the corneal epithelium of rats was removed under anesthesia, followed by the application of a solution containing ruthenium and sodium persulfate (SPS). The corneas were then exposed to blue light at 430 nm at 3 mW/cm^2^ for 5 minutes. Rat corneas were examined and evaluated for corneal opacity, corneal and limbal neovascularization, and corneal epithelial regeneration at days 0, 1, 3, 6, 8, and 14. On day 28, the corneas were isolated for subsequent tissue follow-up and analysis. CXL with ruthenium and blue light showed rapid epithelial healing, with 100% regeneration of the corneal epithelium and no corneal opacity by day 6. The ruthenium group also exhibited significantly reduced corneal (p<0.01) and limbal neovascularization (p<0.001). Histological analysis revealed no signs of cellular damage or apoptosis, which further confirms the biocompatibility and nontoxicity of our method. Confocal and scanning electron microscopy (SEM) images showed a greater density of collagen fibrils, indicating efficient crosslinking and enhanced structural integrity. This study confirmed the in vivo safety, biocompatibility, and functionality of ruthenium and blue light CXL. This method can prevent toxicity caused by UV-A light and can be a rapid alternative treatment to standard crosslinking procedures.

**Graphical Abstract:** 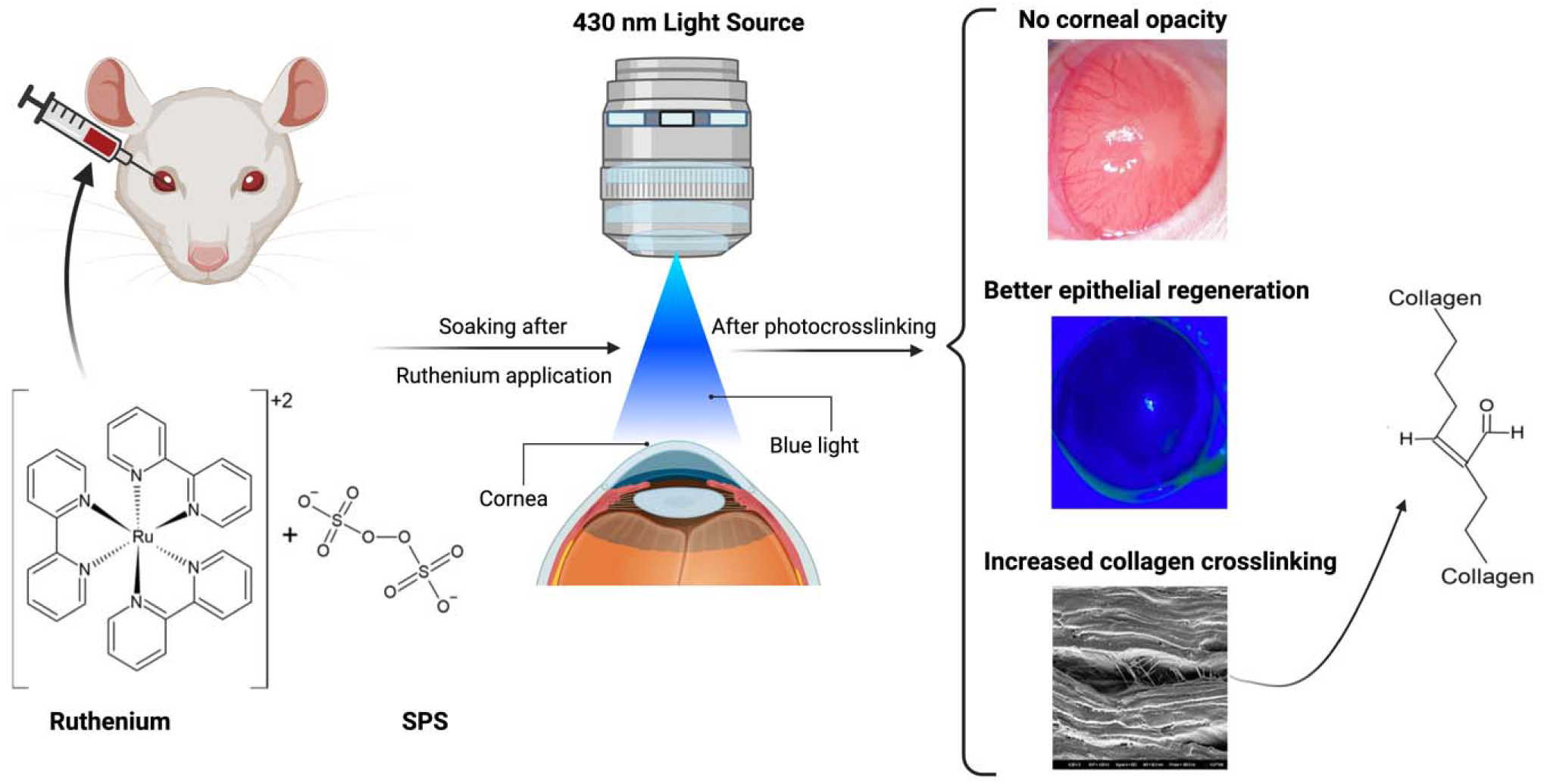

## 1. Introduction

The cornea is a clear avascular ocular tissue that serves as a protective barrier against infections and [1, 2] is responsible for most of the refractive power of the eye [3]. However, corneal thinning diseases distort vision, affecting daily life activities and the mental health of patients [4]. Keratoconus is a corneal disease characterized by bulging of the corneal surface due to progressive weakening and thinning of the corneal stroma [5–7]. A clinically proven method for treating patients with keratoconus is corneal crosslinking (CXL) [8]. This process utilizes riboflavin as a photosensitizer and ultraviolet A (UV-A) light to generate reactive oxygen species (ROS). These ROS induce the formation of covalent bonds between collagen molecules and fibrils, thereby increasing biomechanical stiffness and rigidity, thus preventing further thinning of the cornea [9]. Although commonly used as a nonsurgical treatment option for keratoconus in clinical settings, this method has significant limitations and complications [10]. Notably, the standard riboflavin CXL protocol is not feasible for patients whose corneal thickness is less than 400 µm [11]. When applied to thinner corneas, the UV-A beam reaches corneal endothelial cells, causing endothelial toxicity [12–14], permanent corneal hazing [15], keratocyte toxicity [16] and persistent corneal edema [17], which further requires penetrating keratoplasty [18]. Additionally, adherence to the rule of 400 µm is clinically limiting because patients with advanced-stage ectasia often have corneas thinner than 400 μm [19]. Modifications are made to the standard Dresden CXL procedure to improve its efficacy and reduce patient discomfort [20–28]. Despite these modifications to standard CXL procedures, none have significantly improved the outcomes of CXL treatment. Therefore, there is a need for a new photoinitiator that can achieve corneal collagen crosslinking without causing any ocular toxicity and has excellent corneal penetration capabilities.

We recently introduced a novel photocrosslinking technique utilizing ruthenium and visible blue light as a promising alternative approach for cornea crosslinking [29]. Ruthenium is a water-based photoinitiator that utilizes visible blue light between 400-450 nm to covalently crosslink free tyrosine groups on collagen chains [30]. Ruthenium (Ru(bpy)3) in the presence of sodium persulfate (SPS) and blue light, is oxidized from Ru^2+^ to Ru^3+^ by donating electrons to SPS. After accepting electrons, SPS dissociates into sulfate anions and radicals. These radicals form new covalent bonds between nearby tyrosine groups on collagen chains [31, 32]. Ruthenium and blue light CXL proved to be safe for corneal epithelial cells, limbal stem cells, and fibroblast cells in vitro. In *ex vivo* bovine corneas, crosslinking with ruthenium and blue light resulted in significantly greater corneal stiffness, resistance to enzymatic digestion, and improved optical properties compared to the riboflavin CXL technique [29].

The current study is an extension of our previous work aimed at comprehending the biocompatibility and effectiveness of the ruthenium/ blue light CXL approach in vivo. To achieve this aim, we performed CXL on rat corneas with either ruthenium/blue light or riboflavin/UV-A and compared the two CXL procedures. Our results confirmed the nontoxicity and biocompatibility of ruthenium and blue light. We also showed that ruthenium preserves the integrity and organization of collagen fibers in the corneal stroma while also promoting enhanced bond formation between collagen fibrils.

## 2. Materials & Methods

### 2.1. Animal Ethics

The Institutional Animal Care and Use Committees of Koç University approved all experimental procedures (Approval No: 2019.HADYEK.035) in accordance with Directive 2010/63/EU of the European Parliament and of the Council on the Protection of Animals Used for Scientific Purposes and the Republic of Turkey 5199 Animal Protection Law, Article 9 and 17

### 2.2. Animal Preparation and CXL Procedure

Eight-week-old male Wistar Albino rats were subjected to the CXL procedures. Each group included 4-6 animals as control, ruthenium-CXL, and riboflavin-CXL. Procedures underwent the right eyes, and the left eyes were used as an individual control to check for any complications. The animals were given an intraperitoneal injection of xylazine (5 mg/kg) and ketamine (33 mg/kg) for anesthesia. The depth of anesthetic was confirmed by toe pinch, tail pinch, and corneal reflex tests. For CXL application, the corneal epithelium was removed with sterile disposable blades 7 mm in diameter under topical local anesthesia with 0.5% Alcaine eye drop, and de-epithelization was checked with fluorescein staining under an ophthalmic microscope. The standard riboflavin and UV-A crosslinking procedure includes a UV-A dose of 5.4 J/cm2 in patients with corneal thicknesses greater than 400 μm. Since the corneal thickness of Wistar albino rats is approximately 159 μm [33], low UV-A doses of 1.8 J/cm^2^ (3 mW/cm^2^ for 10 minutes) and 0.9 J/cm^2^ (3 mW/cm^2^ for 5 minutes) were used for the crosslinking procedure. To initiate riboflavin crosslinking, the cornea was topically treated with a 0.1% riboflavin solution (MedioCROSS D, Avedro, Waltham, MA) at regular intervals of 5 minutes for 30 minutes. Subsequently, 365 nm UV-A light illumination was applied at 3 mW/cm2 to activate the crosslinking process. Following the completion of the procedure, a four-times-daily regimen of Tobradex, comprising dexamethasone and tobramycin (Tobradex; Alcon Laboratories Barcelona, Spain), was instilled into the corneas of the animals to prevent ocular infections.

To protect peripheral regions of the eye and minimize complications arising from excessive light exposure during the procedure, a metal ring was placed on the rat cornea, as shown in (Figure 1). This ring served as a protective barrier, covering the limbus area from excessive UV-A irradiation.

**Figure 1.**
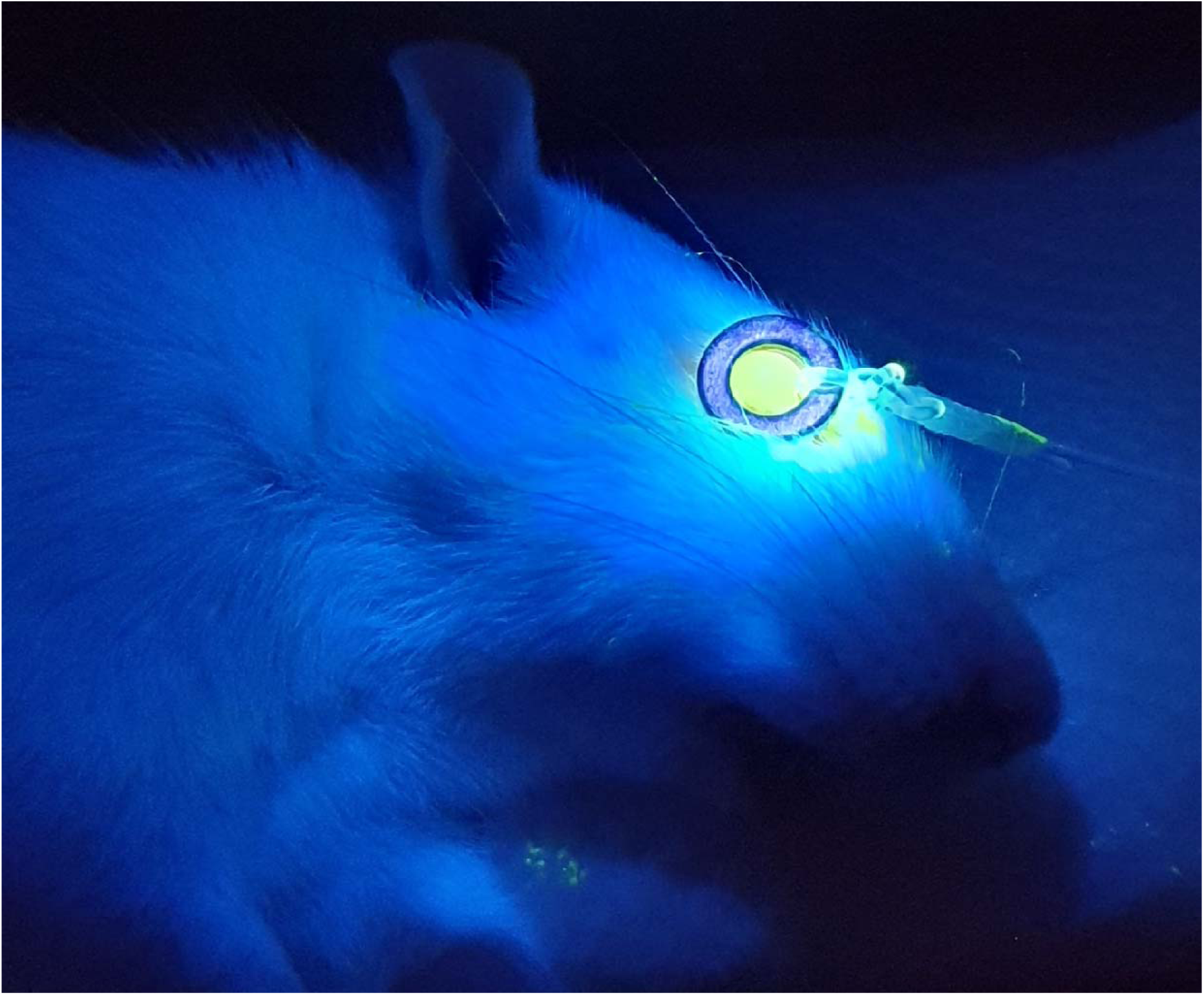
A metal ring with an opening of 7 mm in diameter was positioned on the ocular surface to limit UV-A irradiation to the peripheral cornea.

In the ruthenium group, the de-epithelization process was carried out similarly to that in the riboflavin group. Then, a solution of 1 mM ruthenium and SPS was carefully dropped onto the cornea, followed by a 10-minute incubation period. In our previous study, using ex vivo bovine corneas we demonstrated the efficient crosslinking capability of ruthenium within 5 minutes of blue light exposure. Therefore, we exposed the corneas of the rats to 430 nm blue light at 3 mW/cm^2^ for 5 minutes to initiate the crosslinking process. After the crosslinking procedure, Tobradex eye drops were applied four times daily until the corneal epithelium healed.

### 2.3. Clinical examination

The opacity, neovascularization, and wound healing of the treated corneas were examined under an ophthalmic microscope and photographed under general anesthesia (3% induction; 2% isoflurane during examination) and topical anesthesia (Alcaine) on days 1, 3, 6, 8 and 14 after the operation.

#### 2.3.1. Corneal opacity scoring

The scoring criteria were as follows:

0 = No opacity
1 = Minimal superficial opacity
2 = Slight (stromal) opacity with visible pupillary margin and iris vessels
3 = Middle stromal opacity with only the pupillary margin visible
4 = Dense stromal opacity with anterior chamber
5 = Maximum corneal opacity that completely covers the anterior chamber

#### 2.3.2. Corneal neovascularization scoring

Neovascularization refers to the development of a neovascular branch extending from the corneoscleral limbus to the central cornea. The assessment of neovascularization involved dividing the cornea into quadrants and measuring the extent of neovascularization in each quadrant. Neovascularization was graded per corneal quadrant according to the extent of vascular invasion toward the central cornea [34]. Subsequently, the scores obtained for each quadrant were added to calculate the mean corneal neovascularization score.

The criteria used for scoring were as follows:

0 = No corneal neovascularization
1 = Mild corneal neovascularization
2 = Moderate corneal neovascularization
3 = Severe corneal neovascularization

#### 2.3.3 Limbal neovascularization scoring

The thickening of the vessels around the corneoscleral limbus due to the applied treatment was evaluated under an ophthalmic microscope using a scoring system ranging from 0 to 3.

The scoring criteria were as follows:

0 = Normal limbus neovascularization
1 = Mild limbus neovascularization
2 = Moderate limbus neovascularization
3 = Severe limbus neovascularization

#### 2.3.4 Corneal Epithelial Damage Assessment

The postoperative healing process of the corneal epithelium in rats was monitored by applying fluorescein stains at 1, 3, 5, 7, and 14 days following the CXL procedure. Briefly, the fluorescein strip was placed in 3 ml of saline solution to prepare the dye solution. Green fluorescein staining was applied, and the healing of the epithelial layer was examined with cobalt blue light and captured through photography. The regions retaining fluorescein dye indicate incomplete healing of the epithelial layer. The areas with unsealed epithelium were manually measured using a ruler.

To assess the wound healing progress, the closure diameters were calculated, and the wound healing rate was calculated as a percentage based on the closure measurements.

### 2.4. Corneal Tissue Examination and Histological Analysis

Following the surgical procedure, the condition of the corneas was followed for 28 days. On the 28th day, the rats were euthanized under deep anesthesia, and their eyes were enucleated for subsequent tissue follow-up and analysis. For this procedure, tissues were fixed in freshly prepared 4% paraformaldehyde (PFA) for 1 day at +4°C. Afterward, the tissues were delicately preserved in solutions of 10%, 20%, and 30% sucrose, ensuring gradual stabilization. Next, the tissue specimens were carefully embedded in a cryomold, surrounded by OCT freezing medium, and promptly frozen on dry ice, preserving their structural integrity for further analysis. The frozen tissues were stored at -80°C for an extended duration prior to sectioning. 10 μm -thick sections were taken from the tissues with a cryostat device (Leica CM 1950) and transferred to Superfrost Plus adhesion microscope slides (Epredia, Breda, Netherlands). For histopathological examination, cornea sections were stained with hematoxylin and eosin and observed under a light microscope (DMiL, Leica). Briefly, H&E staining was performed on dehydrated tissue sections by first dipping them in PBS and then incubating the sections in a hematoxylin solution for 2 minutes. After the samples were rinsed with water to remove excess dye, the slides were placed in a jar filled with eosin solution 5 times. Slides were washed with water and dehydrated through a graded alcohol series of 50%, 70%, 90%, and 100% EtOH. Finally, the sections were cleared in xylene and mounted with Entellan medium.

### 2.5. Corneal cell viability by TUNEL assay

Corneal sections (10 μm) were subjected to secondary fixation using 4% PFA in PBS for 20 minutes at room temperature. After thorough rinsing with PBS, the sections were permeabilized with a solution containing 0.1% Triton X-100 in PBS for 2 minutes. After washing with PBS for 5 minutes, the TUNEL assay (In Situ Cell Death Detection Kit, Roche Diagnostics, San Francisco, CA) was performed according to the protocol specified by the manufacturer. Briefly, 50 µL of the DNA Labeling Solution in the TUNEL Assay Kit - BrdU-Red (ab66110, Abcam, MA, USA) was added to the sections, which were then incubated for 1 h at 37°C in the dark. After incubation, the slides were washed with PBS for 10 minutes and treated with BrdU-Red antibody solution for 30 minutes at room temperature. The slides were further washed for 5 minutes with PBS to remove any remaining antibody. Mounting media containing DAPI was applied, and the coverslips were sealed to secure the sections in place. Negative controls were prepared similarly but without the use of an antibody solution. All immunofluorescence images were taken using an upright fluorescence microscope (Axio Imager M1, ZEISS, Oberkochen, Germany).

### 2.6. Protein and collagen expression in corneal tissue

Protein and collagen expression in the corneas of the experimental groups was analyzed via immunofluorescence staining. Dehydrated tissue sections were incubated with 0.1% Triton X-100 in PBS for 8 minutes for permeabilization. Then, they were incubated with Superblock (Thermo Fisher Scientific, MA, USA) for 10 minutes at room temperature for blocking. After washing with PBS, the samples were treated with rabbit anti-CK12 primary antibody (MA5-42701, Thermo Fisher Scientific, MA, USA), mouse anti-CD31 primary antibody (ab24590, Abcam, MA, USA) and rabbit anti-collagen I primary antibody (Pa1-26204, Thermo Fisher Scientific, MA, USA) at 1:100 dilutions in blocking solution overnight at 4°C. After three washes with PBS, the samples were incubated with Alexa Flour 532-conjugated anti-mouse secondary antibody at a 1:200 dilution (A-11002, Invitrogen, MA, USA) and Alexa Flour 488-conjugated anti-rabbit secondary antibody at a 1:200 dilution (A-11008, Invitrogen, MA, USA) for 90 minutes at 37°C. After washing with PBS, the samples were mounted with 406-diamidino-2-phenylindole (DAPI, ab104137, Abcam, MA, USA) mounting media. Negative controls were stained similarly without the addition of primary antibodies. Anti-CK12 and anti-CD31 immunofluorescence images were acquired by using an immunofluorescence inverse light microscope (Axio Observer Z1, Zeiss, Oberkochen, Germany), and anti-collagen I immunofluorescence images were taken by confocal laser scanning microscopy (DMI8 SP8, Leica, Wetzlar, Germany).

### 2.7. Scanning electron microscopy (SEM)

Following the CXL procedures, cornea samples from all groups were collected for SEM analysis. SEM images of the samples were taken using a Zeiss EVO LS 15 SEM. Prior to SEM imaging, 10 μm cryosectioned cornea samples were carefully mounted on metal stubs using carbon adhesive tape. Subsequently, a thin layer of approximately 20 nm gold coating was applied to the samples through sputtering using a 108 Auto Sputter Coater with an MTM-10 Thickness Monitor (Cressington, Watford, UK).

### 2.7. Statistical analysis

The statistical analysis was performed with GraphPad Prism software (GraphPad Software Inc., San Diego, CA, USA). The data are presented as the mean ± SEM. Comparisons between the experimental data of groups were performed using two-way ANOVA or one-way ANOVA with Tukey’s comparison test, where a p-value less than 0.05 was considered significant for all experiments. The standard deviation of the mean is indicated by error bars for each data group.

## 3. Results

In this study, we performed corneal crosslinking to reinforce the cornea by creating additional crosslinks between collagen fibers in Wistar albino rats. The corneas were treated with ruthenium/blue light and compared with both riboflavin/UV-A treated and control corneas to assess the effectiveness and safety of our proposed CXL procedure in vivo. The crosslinking setup is shown in Supp. Figure 1.

### 3.1. In vivo corneal crosslinking and scoring

Ensuring proper healing of the cornea is crucial to avoid excessive opacity and the development of corneal and limbal neovascularization, which can impede light transmission. Microscopy images of the cornea on days 0, 1, 3, 6, 8, and 14 are shown in (Figure 2A), providing insights into the healing process. For corneas crosslinked with ruthenium and blue light, the mean corneal opacity was 0.1±0.030 on day 1 and increased to 0.6±0.050 on day 3, as shown in (Figure 2B). No corneal opacity was observed in the ruthenium group on day 6 or later.

**Figure 2.**
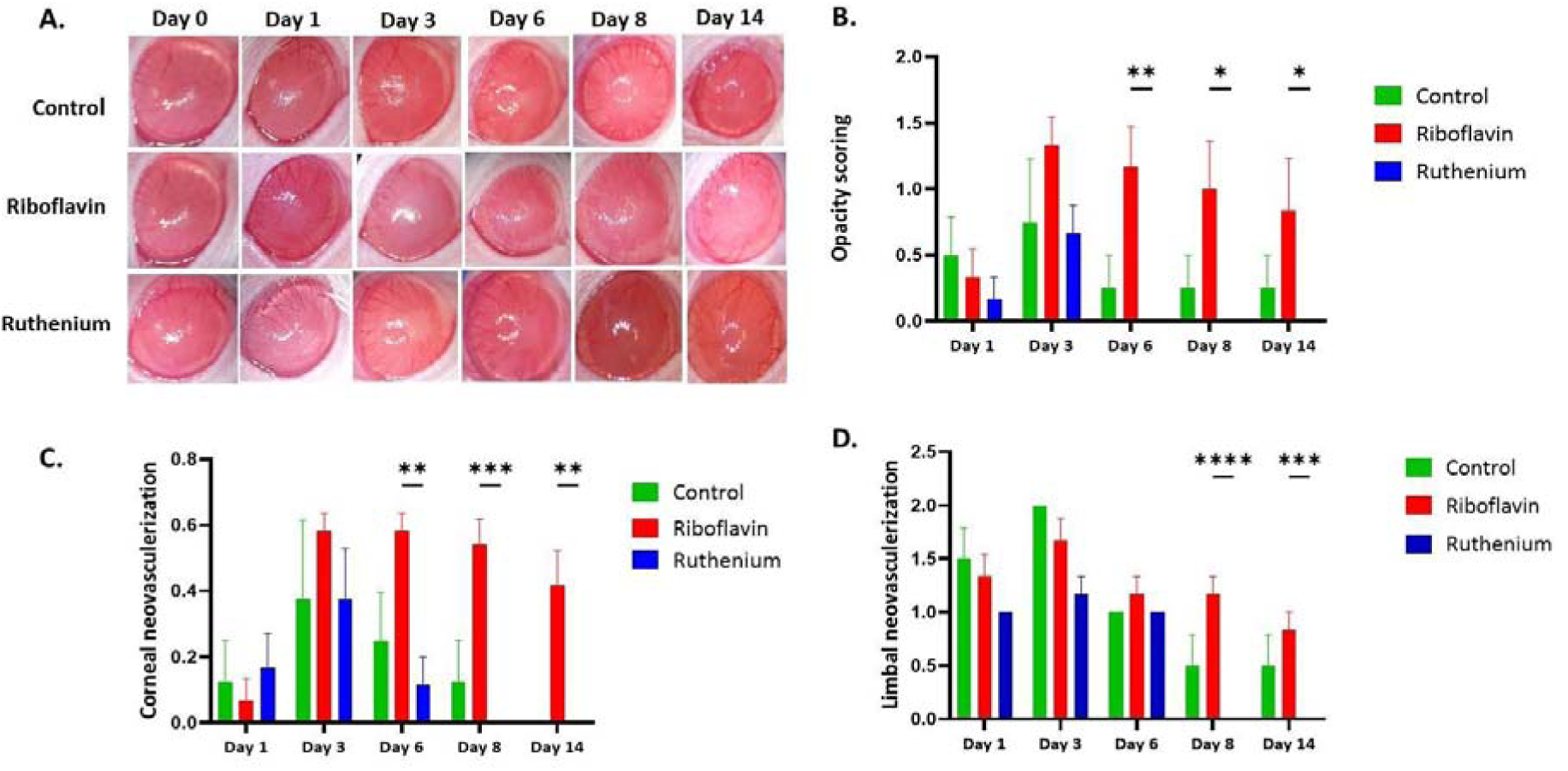
Corneal examination after the crosslinking procedure. (A) Eye images of the groups were taken under an ophthalmic microscope at different time intervals after the crosslinking procedure. Quantitative analysis of the scores calculated for each time interval. (B) Corneal opacity increased in all groups until day 3. The ruthenium group had a clear cornea from day 6 to day 14. Despite the improvement observed in the riboflavin group after day 3, there was still a noticeable and statistically significant difference in corneal opacity between the ruthenium-treated group and the riboflavin-treated group until day 14. (C) There was less corneal neovascularization in the ruthenium group than in the riboflavin group after day 6. Compared with the riboflavin group, the ruthenium group showed no corneal neovascularization, whereas the riboflavin group exhibited significantly greater corneal neovascularization scores on days 8 and 14. (D) The limbal neovascularization score showed that the ruthenium group had low vascularity in the cornea, and this effect was not detected after day 8. The data are expressed as the mean ± SEM (n = 4-6 rats/group), * p < 0.05, ** p < 0.01, *** p < 0.001, **** p < 0.0001).

We observed severe complications in the riboflavin group exposed to a UV-A dose of 1.8 J/cm2. These complications include corneal perforation, bulging of the cornea outside and plaque formation, as shown in (Supp. Figure 2). To mitigate this undesirable outcome, we further reduced the UV-A dose from 1.8 J/cm2 to 0.9 J/cm2. This modification led to a more favorable outcome in our experimental control group and we continued further experiments with low a UV-A dose. The riboflavin group exposed to 0.9 J/cm2 exhibited a reduced corneal opacity score of 0.5±0.040 on day 1 and 0.75±0.030 on day 3. However, despite the decrease in the corneal opacity score after day 3, there was a significant difference between the ruthenium and riboflavin groups until day 14 (p<0.05), with the ruthenium group displaying a clear cornea without any complications.

Our observation of corneal and limbal neovascularization revealed no significant differences among the three groups until day 3 (Figure 2C). However, on day 6, the corneal neovascularization score for the ruthenium group was 0.11±0.02, whereas that for the riboflavin group was 0.58±0.06. After day 6, the corneas in the ruthenium group showed no signs of corneal neovascularization, whereas those in the riboflavin group exhibited significantly greater corneal neovascularization scores than those in the ruthenium group on days 8 (p<0.003) and 14 (p<0.01). Notably, the difference between the ruthenium group and the control group was not statistically significant (p>0.05). Figure 2D shows the limbal neovascularization scores across all the groups. There was no significant difference between the three groups until day 6. However, on day 8, we observed no limbal neovascularization in the ruthenium group, while the riboflavin group had limbal neovascularization scores of 1.16±0.04 on day 8 and 0.83±0.03 on day 14, with statistically significant values of p < 0.0001 and p < 0.001, respectively.

Figure 3A shows corneal epithelial wound healing for all the groups after the crosslinking procedure on days 1, 3, 6, 8, and 14. The results revealed significant differences between the ruthenium and riboflavin-treated groups on day 1 (p<0.001) and day 3 (p<0.0001), as shown in (Figure 3B). By day 3, the ruthenium-treated group exhibited a remarkable 86% epithelial recovery, whereas the riboflavin-treated group showed a significantly lower rate of epithelial regeneration, with only a 20% recovery observed. On day 14, both the control and ruthenium-treated groups showed complete regeneration of the epithelium, indicating successful wound healing. Conversely, the riboflavin treated group showed diffuse punctate epitheliopathy and scar formation on days 3 and 6. However, by day 14, the epithelium in the riboflavin treated group had fully regenerated.

**Figure 3:**
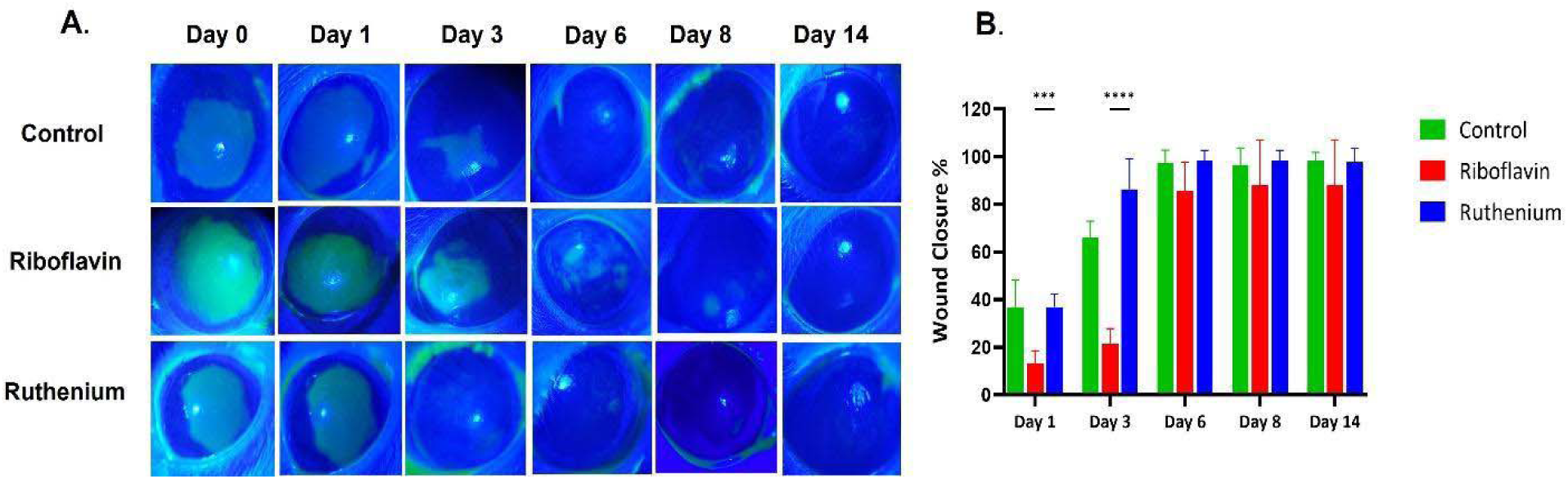
Epithelial wound healing rate with fluorescein staining after corneal crosslinking. (A) Fluorescein-stained images of the control, riboflavin, and ruthenium groups were taken 0, 1, 3, 6, 8, and 14 days after the crosslinking procedure. The green color shows the stained areas, which indicate the absence of epithelial cells on the wounds. (B) The percentage of epithelial wound closure was calculated and compared at each time point. Wound closure in the ruthenium group was greater than that in the riboflavin group on days 1 and 3. The data are expressed as the means ± SEMs (n = 4-6 rats/group, * p < 0.05, ** p < 0.01, *** p < 0.001, **** p < 0.0001).

### 3.2. Tissue examination

After the initial crosslinking procedure, the corneas were evaluated over a period of 28 days. On the 28th day, the rats were euthanized under anesthesia, and their eyes were removed and subjected to further tissue analysis. The histopathological findings obtained from hematoxylin & eosin staining highlighted the differences in corneal structure among the treatment groups (Figure 4A). The results indicated that in the control, riboflavin, and ruthenium groups, the epithelial, stromal, and endothelial layers of the cornea appeared normal. However, in the riboflavin group treated with a higher UV-A dose of 1.8 J/cm^2^, thickening of the central cornea, loss of the endothelial layer, and vascularization of the stroma were observed (Supp. Figure 2).

**Figure 4.**
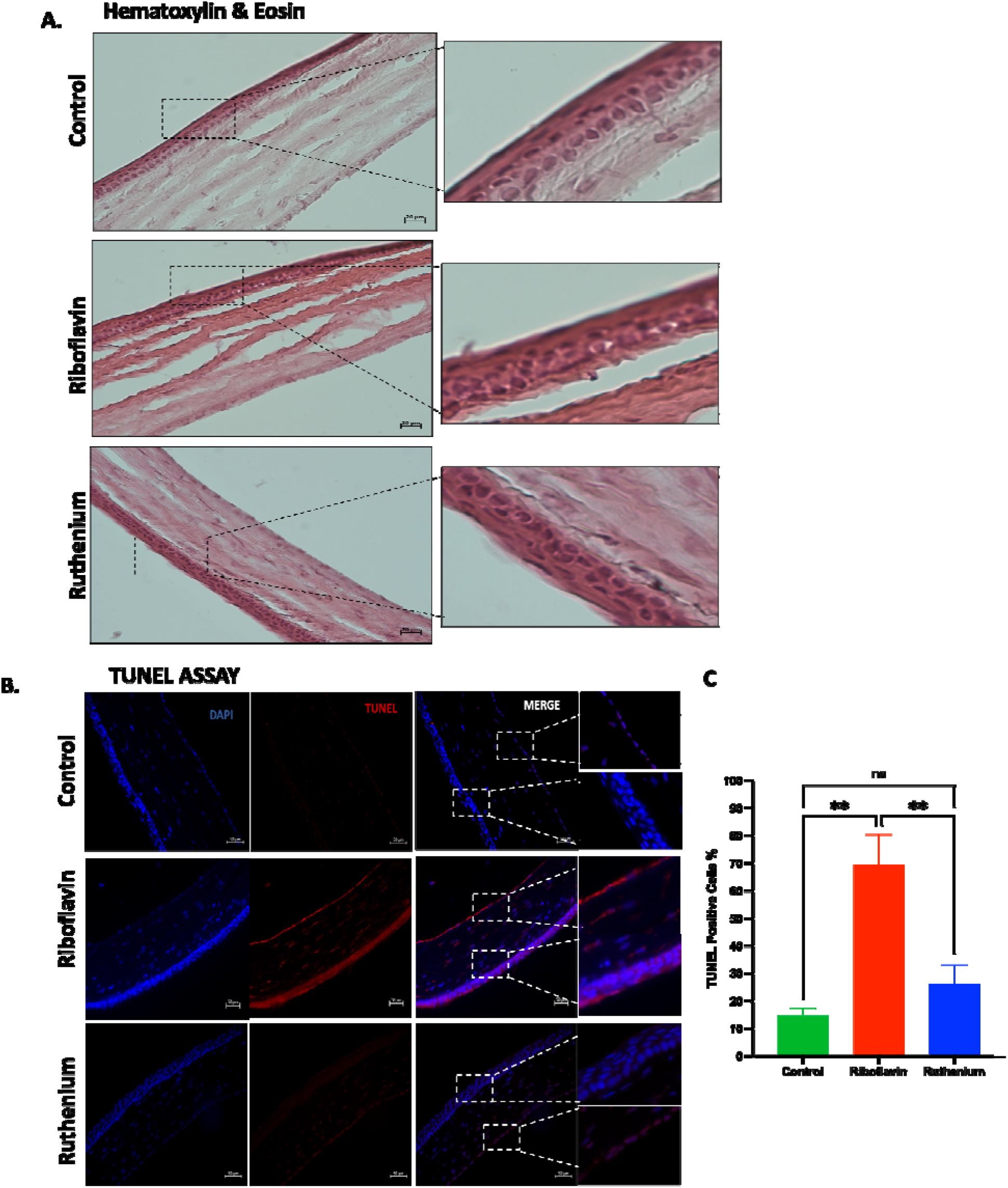
Histological examination of corneal sections. At the end of 28 days, the enucleated eyes were fixed and sectioned for staining. (A) The results of hematoxylin and eosin staining showed that the integrity of the cornea was preserved in all groups. Compared with the control group, the ruthenium group showed no visible alterations in tissue structure. Scale bars represent 20 µm. (B) TUNEL assay indicating corneal crosslinking-induced apoptosis. DAPI (blue) shows cell nuclear immunofluorescence staining, and TUNEL (red) shows DNA fragmentation. (C) TUNEL-positive cells. Compared to those in the control and ruthenium groups, the number of TUNEL-positive cells throughout the cornea in the riboflavin group was greater. However, the number of TUNEL-positive cells in the control and ruthenium groups was not significantly different (p>0.05). Scale bars represent 50 µm.

These observations suggest that the toxic effects of UV-A light cause structural changes and damage to the cornea. On the other hand, the corneal crosslinking method using ruthenium and blue light consistently maintained the normal architecture of all corneal layers. Our findings provide compelling evidence that corneal crosslinking with ruthenium and visible light effectively preserved the integrity of the cornea and prevented the adverse effects observed in the riboflavin group, thus demonstrating its protective effect.

The effect of the crosslinking procedure on keratocyte viability was assessed using terminal deoxynucleotidyl transferase broken-end labeling (TUNEL), which is used to detect DNA fragmentation during cell death. The experimental results revealed that the control group and ruthenium/blue light group exhibited similar numbers of TUNEL-positive cells, as shown in (Figure 4B). This suggests that the presence of ruthenium did not significantly increase cell death compared to that in the untreated control group. However, in the riboflavin/UV-A group, TUNEL-positive cells were distributed throughout the entire cornea, indicating widespread DNA fragmentation and cell death. Moreover, the riboflavin group exposed to a higher UV-A dose of 1.8 J/cm2 exhibited a high rate of apoptotic cell death, which is evidence of the significant impact of UV-A irradiation even 28 days after the initial operation (Supp. Figure 3).

According to the corneal neovascularization scoring results, vascularization at a certain level was observed in the riboflavin group until the end of the experiment. To confirm this finding in tissue sections, the protein expression of CD31 was also examined. CD31 is a widely used marker for blood vessels and provides insights into the dynamics of angiogenesis. Under different conditions, its expression increases in the cornea in response to stimuli. Herein, we examined the expression of CD31 in the experimental groups and detected enhanced signals in riboflavin-treated corneal sections Figure 5A. The presence of CD31 in the central cornea of the riboflavin group supported corneal neovascularization scoring at the tissue level. To assess the wound closure efficiency at the cellular level, we performed Cytokeratin 12 (CK12) staining on the corneas. CK12 is highly expressed in corneal epithelial cells, and under conditions of epithelial disruption, its expression is altered. All groups exhibited similar CK12 expression, consistent with the fluorescein staining results during corneal wound healing (Figure 5 B).

**Figure 5.**
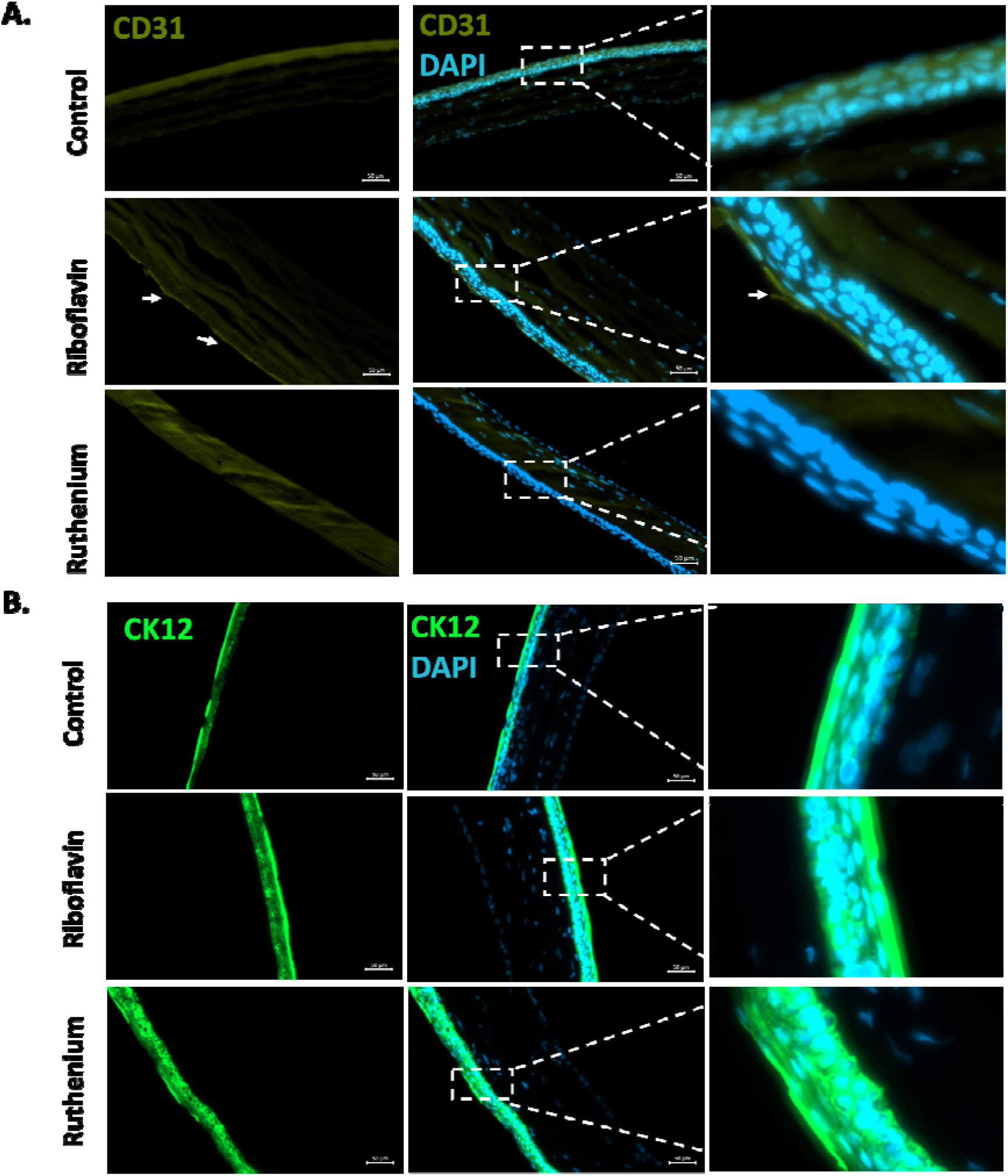
Expression of corneal-specific cell markers in tissue sections. (A) CD31 is used as a marker for angiogenesis and inflammation. In a normal healthy cornea, CD31 expression is not detected. However, during wound healing, corneal neovascularization occurs, and CD31 is expressed in the central cornea. CD31 expression in the Riboflavin group is indicated by white arrows. (B) CK12 is a highly expressed intermediate filament in epithelial cells that maintains corneal structural integrity. All groups had similar CK12-labeled epithelial cells showing wound healing after 28 days of treatment. Scale bars represent 50 µm.

### 3.3. Ultrastructure of corneal collagen fibers

Collagen in the cornea plays a vital role in preserving the structural integrity of corneal tissue. In a healthy corneal stroma, type I collagen is the predominant component of the extracellular matrix [35]. To investigate collagen arrangement in the central cornea, we conducted immunofluorescence analysis using a collagen type-I antibody. Our findings indicate uniform expression of type I collagen in the ruthenium group, closely resembling that of the control and riboflavin groups as illustrated in (Figure 6A). To further examine the structural alterations in corneal tissues and collagen fibers caused by CXL, we conducted cross-sectional imaging of corneal tissues using a scanning electron microscope (SEM). Our observations revealed a notably greater density of collagen fibrils and fewer interstitial spaces between collagen layers in both the ruthenium and riboflavin groups than in the control group, as shown in (Figure 6B). These observations indicate that the ruthenium and blue light CXL led to a similar compact arrangement of collagen fibrils and closer proximity between collagen layers in the corneal tissue to those in the control group.

**Figure 6.**
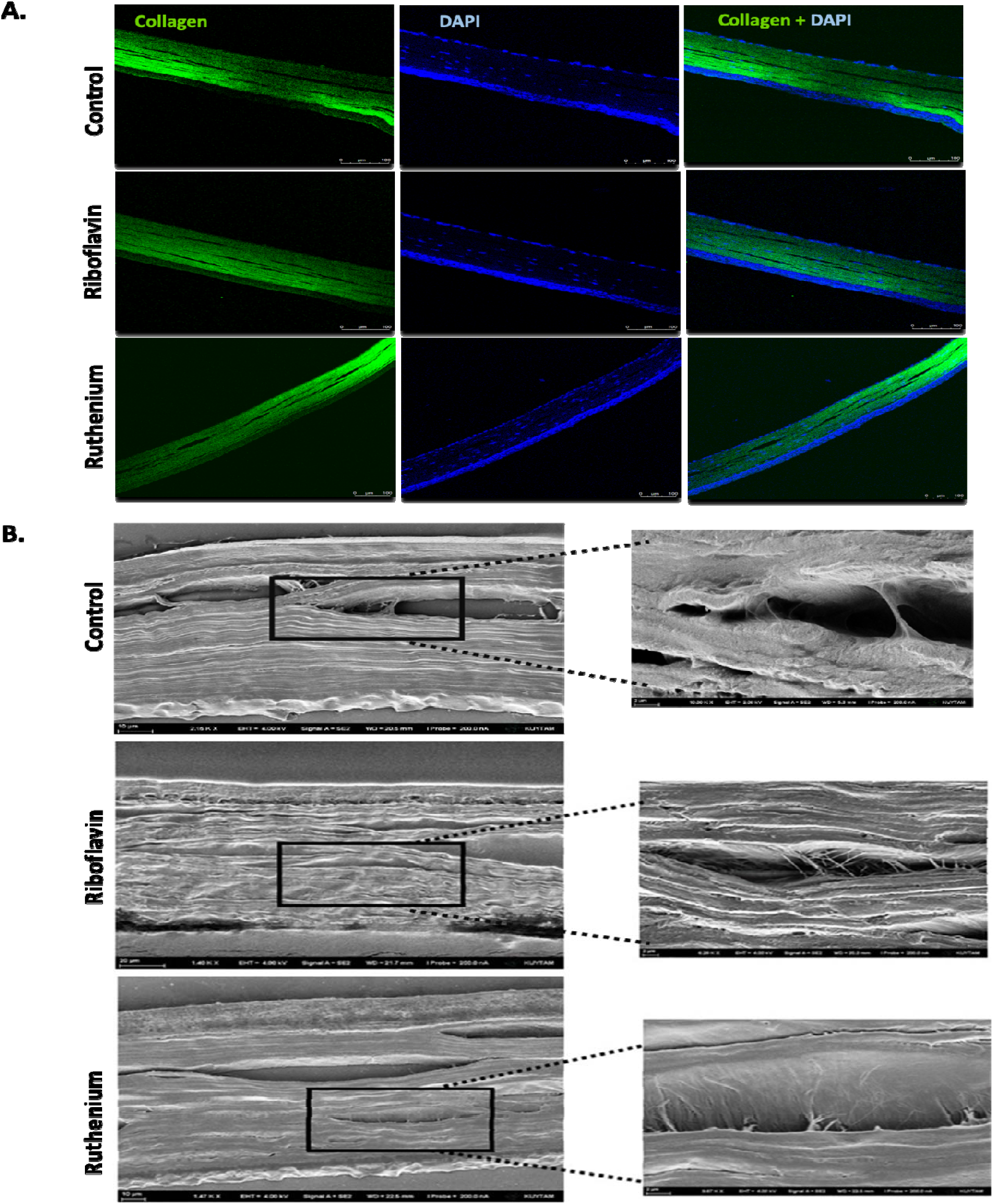
Ultrastructure of corneal collagen fibers. (A) Collagen type-I staining showing the organization of collagen fibers and their integrity in the corneal stroma in all the groups. Scale bars represent 100 µm. (B) Scanning electron microscopy images of the cornea. Both the ruthenium and riboflavin groups exhibited a greater density of collagen fibrils and narrower interstitial spaces between the collagen layers compared to the control group, as shown in the magnified figure (right).

## 4. Discussion

When developing materials for corneal crosslinking, it is paramount to consider ocular cytotoxicity in clinical applications. In the current study, we aimed to investigate the safety and biocompatibility of ruthenium and blue light for corneal crosslinking in vivo. Various parameters, including the presence of opacity in the central cornea, epithelial wound healing, and neovascularization, which indicate inflammation in both the limbus and central cornea, were assessed. It is worth noting that due to the difference in thickness between rat and human corneas (a rat cornea 5x thinner than a human cornea) [36], a modified crosslinking procedure tailored to the characteristics of the rat cornea was applied.

Previous studies have shown that a significant corneal stiffening effect occurs even at a reduced UV-A fluence of 0.09LJJ/cm^2^ [37], while a fluence within the range of 1.62 J/cm^2^ to 2.7LJJ/cm^2^ results in minimal adverse effects such as neovascularization and scar formation [38]. However, our observations showed that a UV-A dose of 1.8 J/cm^2^ resulted in more extensive and irreversible losses in central endothelial cell density. This was accompanied by an increased incidence of perfused eye pathology (Supp. Figure 2), which refers to the development of abnormal blood vessel growth and circulation within the eye. These effects are similar to the described structural changes, scar formation, and keratocyte loss observed in standard CXL procedures performed on humans [39, 40]and rabbits [41]. The control and ruthenium groups showed faster epithelial healing following the crosslinking procedure. Delayed re-epithelialization could be a possible complication of the riboflavin-UV-A-based crosslinking procedure [42]. To mitigate the risk of retinal blue light toxicity, we used a light intensity of 3 mW/cm^2^, which is significantly below the maximum permissible radiant exposure (MPHc) of 54 mW/cm^2^ for wavelengths between 400-450 nm [43]. Our proposed method offers several key benefits, including the use of blue light, which reduces the cytotoxicity, a shorter application time, increased diffusion, and improved crosslinking efficiency.

Given the differences in corneal anatomy and physiological characteristics between rodents and humans, particularly in corneal thickness, optimization experiments are necessary to determine the optimal dosage, light intensity, and treatment duration for crosslinking procedures. Furthermore, additional well-designed studies involving higher organisms, such as rabbits or pigs, are needed to validate the effectiveness and safety of corneal crosslinking with ruthenium-based visible light treatment. Such studies may include comprehensive corneal examination after treatment to measure intraocular pressure levels and corneal curvature posttreatment. Although the ruthenium-based CXL procedure already offers a faster alternative to riboflavin CXL, it is worth exploring the possibility of conducting studies using varying intensities of blue light. This approach has the potential to further decrease the CXL time associated with the ruthenium-based method.

## 5. Conclusion

Based on our findings, the ruthenium and blue light crosslinking procedure did not cause any toxicity in vivo. The tissue integrity and cell viability were well preserved, indicating that the procedure had a favorable impact on the corneal tissue and cells. This can help eliminate the potential complications associated with using UV-A light. Our method can be considered a new rapid, effective, and nonsurgical treatment option for all types of ectatic corneal pathologies, including keratoconus.

## Acknowledgments

This study was supported by the Scientific and Technological Research Council of Turkey (TUBITAK) under the TUBITAK 3501 Program (project number 219S349). The authors gratefully acknowledge using the services and facilities of the Koç University Animal Research Center (KUARF), the Koç University Research Center for Surface Science (KUYTAM), and Koç University Research Center for Translational Medicine (KUTTAM), funded by the Presidency of Turkey, Presidency of Strategy and Budget. The content is solely the authors’ responsibility and does not necessarily represent the official views of the Presidency of Strategy and Budget. The graphical abstract is created with **BioRender.com**

