## Supplementary figures for "A unique and biocompatible approach for corneal collagen crosslinking in vivo"

**Supporting Information**

To ensure accurate measurement and control of the light intensity used during the procedure, we employed a power meter to quantify the intensity of the light being delivered to the cornea. During the crosslinking process, a collimator was utilized to provide focused illumination, directing the light into a narrow beam. In this case, the collimator was positioned above the cornea, allowing the light to be precisely directed onto the treatment area.

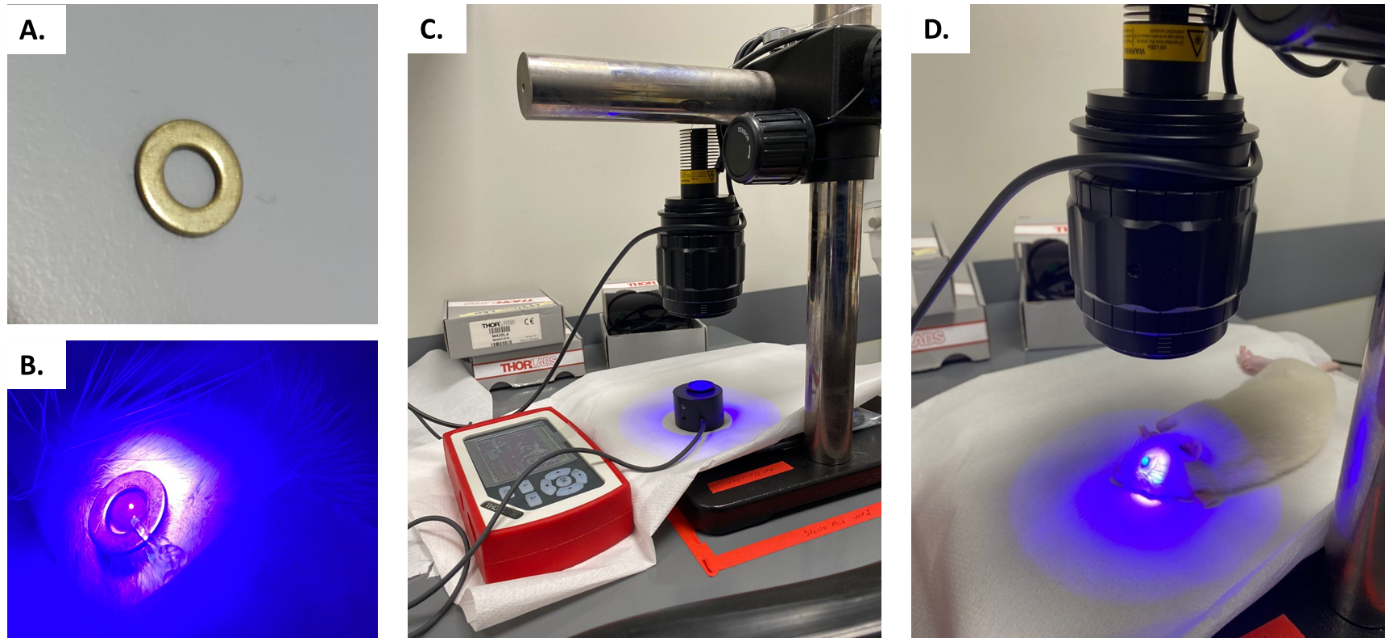

**Supp. Figure 1.** Overview of corneal crosslinking setup. (A) Metal ring 7 mm prepared to protect the limbus, the sclera, and conjunctiva, (B) Position of the ring on the eye during light illumination, (C) Irradiation dose in the lightning setup was measured with a power meter to ensure the correct amount of light given, (D) Application of light illumination on cornea to accelerate crosslinking procedure.

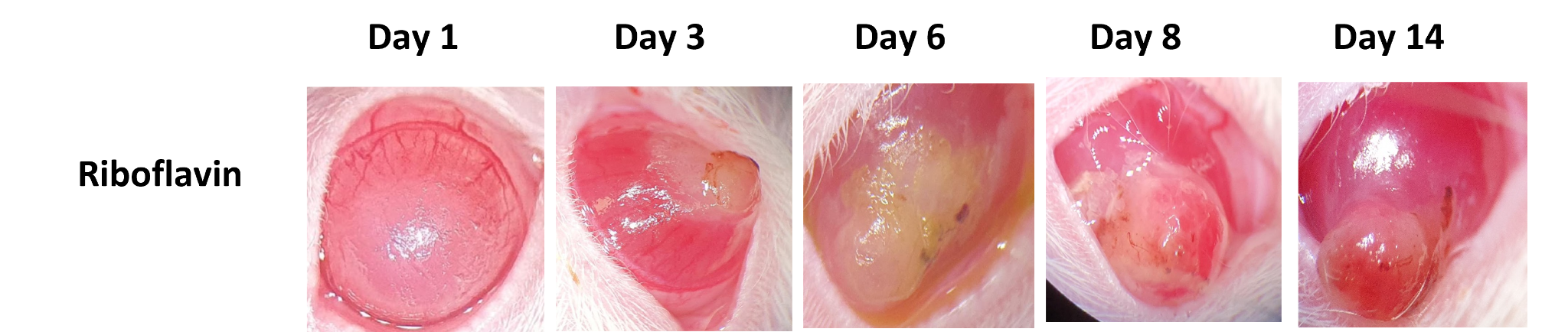

**Supp. Figure 2:** Macroscopic image of the riboflavin group treated with 1.8 J/cm2 UV-A (n=6). Severe complication with corneal perforation observed due to UV-A irradiation in rat corneas. Eyes were treated with antibiotics until the rats were euthanized.

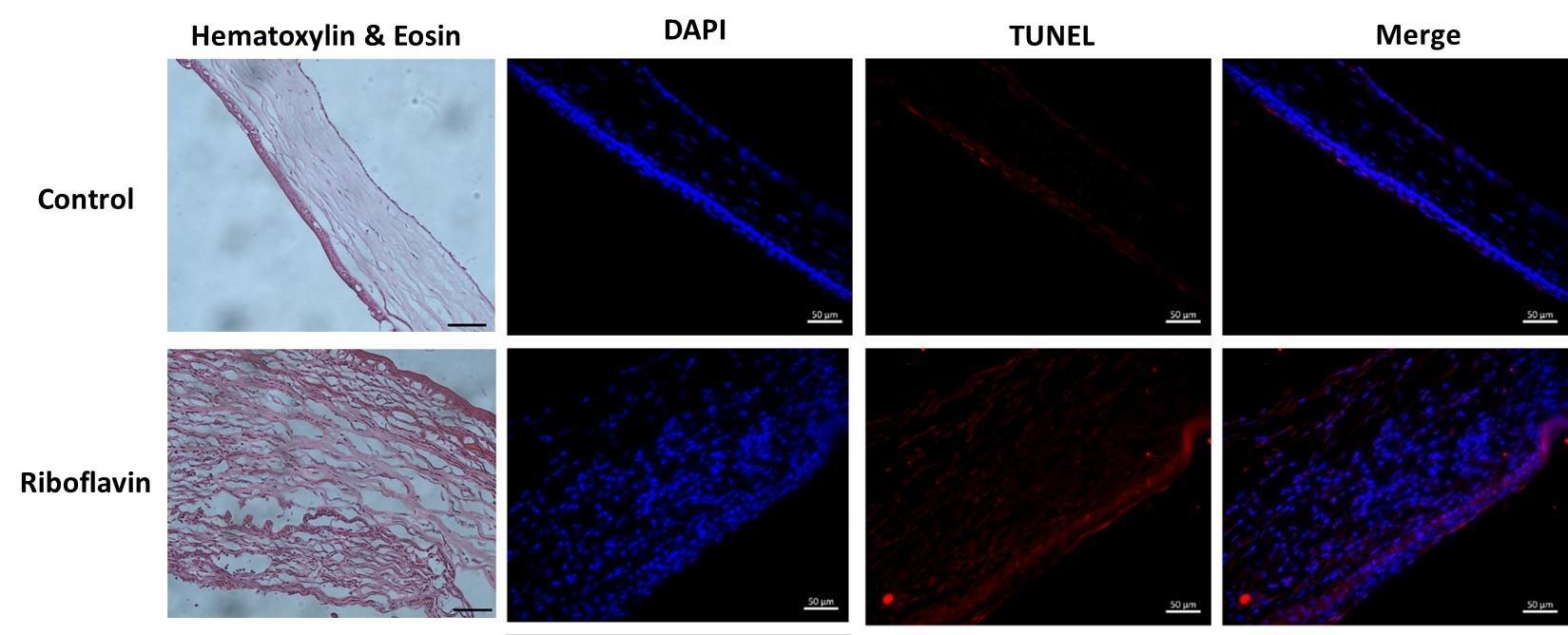
**Supp. Figure 3:** Histological and cell death examination of control and the riboflavin group treated with 1.8 J/cm2 UV-A. Corneal fibrosis was observed as a response to injury in the riboflavin group. Irregular collagen fiber and neovascularization were also found with hematoxylin and eosin staining. Increase in the cell death following 1.8 J/cm2 UV-A. Scale bars represent 50 µm.
